# Temporal modification of H3K9/14ac and H3K4me3 histone marks mediates mechano-responsive gene expression during the accommodation process in poplar

**DOI:** 10.1101/2023.02.12.526104

**Authors:** Ritesh Ghosh, Juliette Roué, Jérôme Franchel, Amit Paul, Nathalie Leblanc-Fournier

## Abstract

Plants can attenuate their molecular response to repetitive mechanical stimulation as a function of their mechanical history. For instance, a single bending of stem is sufficient to attenuate the gene expression in poplar plants to the subsequent mechanical stimulation, and the state of desensitization can last for several days. The role of histone modifications in memory gene expression and modulating plant response to abiotic or biotic signals is well known. However, such information is still lacking to explain the attenuated expression pattern of mechano-responsive genes in plants under repetitive stimulation. Using poplar as a model plant in this study, we first measured the global level of H3K9/14ac and H3K4me3 marks in the bent stem. The result shows that a single mild bending of the stem for 6 seconds is sufficient to alter the global level of the H3K9/14ac mark in poplar, highlighting the fact that plants are extremely sensitive to mechanical signals. Next, we analyzed the temporal dynamics of these two active histone marks at attenuated (*PtaZFP2, PtaXET6*, and *PtaACA13*) and non-attenuated (*PtaHRD*) mechano-responsive loci during the desensitization and resensitization phases. Enrichment of H3K9/14ac and H3K4me3 in the regulatory region of attenuated genes correlates well with their transient expression pattern after the first bending. Moreover, the levels of H3K4me3 correlate well with their expression pattern after the second bending at desensitization (3 days after the first bending) as well as resensitization (5 days after the first bending) phases. On the other hand, H3K9/14ac status correlates only with their attenuated expression pattern at the desensitization phase. The expression efficiency of the attenuated genes was restored after the second bending in the histone deacetylase inhibitor-treated plants. While both histone modifications contribute to the expression of attenuated genes, mechanostimulated expression of the non-attenuated *PtaHRD* gene seems to be H3K4me3 dependent.

## INTRODUCTION

Plants have sophisticated molecular machinery to perceive various mechanical cues, like wind, rain, touch, and bending. Exposure to repeated mechanical stimulation over a period leads to severe alterations in plant growth and development, referred to as thigmomorphogenesis (Jaffe, 1973). For instance, repetitive bending of stem (to mimic wind stimulation) causes various thigmomorphogenic responses in trees, like reduction in longitudinal growth, increase in diameter growth, and modification of wood’s chemical and mechanical properties (Telewski and Pruyn, 1998; Pruyn *et al*., 2000; Kern *et al*., 2005; Roignant *et al*., 2018). So far, mechano-stimulated molecular responses in plants have been mostly studied after single or multiple simultaneous touch treatments without intervals, which did not take account of natural conditions, where stems and foliage constantly encounter repetitive mechanical stimulation at different frequencies. As an example, wind gusts cause repeated flexing of plant organs at different frequencies in the range of 1–5 Hz, corresponding to 60–300 bends per minute (Rodriguez *et al*., 2008). Moreover, several studies indicate that plant responses to repetitive mechanostimulation are more complex as it depends on treatment intensity and frequency (Ghosh *et al*., 2021). To understand the impact of repetitive mechanical stimulation on a tree, Martin *et al*. (2010) measured daily diameter growth variation of poplar plants after quantitative repeated bending of stems. Interestingly, an attenuated diameter growth response was observed after the second bending relative to the first one when two treatments were applied with an interval of 4 days. The same study also showed that an interval of 10 days is necessary between two bending treatments to recover the diameter growth response. These data demonstrated that a single bending is sufficient to alter the plant responsiveness to mechanical cues, and the state of desensitization can last for several days. Corroboratively, a transcriptomic analysis showed that the expression of a large set of mechano-responsive genes can be attenuated in poplar stems when two bending treatments are separated by 24 h intervals (Pomies *et al*., 2017). Based on this transcriptomic analysis, 96% of poplar mechano-responsive genes were transiently expressed after a single bending but showed an attenuated response by the second bending. This analysis revealed another category of genes – non-attenuated genes – which show a similar expression pattern after the first or the second bending (Pomies *et al*., 2017). Such desensitization of mechanoresponse was later confirmed in Arabidopsis, where repeated touch treatment within 24 h gradually attenuates the expression of a large set of mechano-responsive genes, including touch marker genes, like *WRKY15* and *WRKY40* (Xu *et al*., 2019). Besides gene expression, attenuated Ca^2+^ spiking and eATP release in Arabidopsis roots have also been noted in response to repetitive mechanostimulation (Legue *et al*., 1997; Weerasinghe *et al*., 2009). Collectively, these results indicate that the acclimation of responses to repeated mechanostimulation has a global impact on the cellular process. Such acclimation of mechanosensitivity is known as the ‘‘accommodation’’ phenomenon (Leblanc-Fournier *et al*., 2014). Intuitively, such self-regulation could be an adaptive strategy to fine-tune plant responses to repetitive events like wind by preventing over-reaction and costly investment in reducing or redirecting growth. For instance, beech trees respond only to the intense wind but not to chronic lower-intensity wind signals (Bonnesoeur *et al*., 2016).

Plants can remember past events and modulate their responses to subsequent stimulation accordingly. Multiple studies show that exposure to biotic and abiotic stressors can generate molecular memory in plants via epigenetic modifications (Baurle and Trindade, 2020). Epigenetic modifications alter the accessibility of genes to transcription regulatory proteins without affecting the DNA sequence. By providing a long-term somatic memory at the level of gene expression, modification of amino acid residues of histone N-terminal tails seems to contribute to plants’ adaptation to various environmental stresses. For instance, the deposition of H3K4me3 transcription activation mark was observed at the promoters of heat shock and dehydration memory genes which could be an important strategy to mark recently transcribed genes for stronger and faster reactivation upon exposure to recurrent stress (Ding *et al*., 2012; Lamke *et al*., 2016). It has been shown that histone modifications facilitate the defense priming process (Conrath *et al*., 2015). For example, repetitive exposure to various abiotic stresses primes the expression of pattern-triggered immunity (PTI) genes through the enrichment of acetylation of histone H3 lysine 9/14 (H3K9/14ac) and methylation of histone H3 lysine 4 (H3K4me2 and H3K4me3) at their promoter and increases bacterial resistance (Singh *et al*., 2014). Contrary to the attenuated expression of genes by repetitive mechanical stimulation, primed plants show faster or stronger reactivation of stress memory genes under recurrent biotic or abiotic stress.

However, the molecular mechanism which alters the plant responsiveness to repetitive mechanical cues in plants is yet to be elucidated. A couple of preliminary studies performed on Arabidopsis indicate the involvement of epigenetic regulation during plant responses to mechanical stimulation. For example, using Arabidopsis mutant, Cazzonelli *et al*. (2014) showed that histone lysine methyltransferase SDG8 regulates the expression of 8.7% of the touch-responsive genes. This enzyme belongs to the Trithorax-group (TrxG) proteins, which catalyze H3K4 and H3K36 methylation. The same study also reported a reduced H3K4me3 level at the *TCH3* locus in the *sdg8* mutants, which correlates well with its expression pattern. VIP3 encodes part of the RNA polymerase II-associated factor 1 complex Paf1, which is required for H3K36me3 active mark at *TCH3* and *TCH4* loci and their mechano-stimulated expression (Jensen *et al*., 2017). Interestingly, both *sdg8* and *vip3* mutants are impaired in thigmomorphogenesis (Cazzonelli *et al*., 2014; Jensen *et al*., 2017). However, none of these studies characterized how histone marks change with time after mechanostimulation and whether they are involved in the desensitization-resensitization process during repetitive stimulation. Histone modifications can be reversible to activate or deactivate gene expression (Justin *et al*., 2010; Berr *et al*., 2011). Thus, we hypothesized that the kinetics of histone modifications between one and two mechanical stimulations at the regulatory regions of mechano-responsive genes could be an additional layer to control the plant responsiveness to repetitive mechanical cues. Using poplar as a model plant in this study, we investigated the temporal dynamics of H3K9/14ac and H3K4me3 active marks in the mechanostimulated poplar stem and their contribution to the expression of attenuated and non-attenuated genes during the desensitization-resensitization phases.

## MATERIALS AND METHODS

### Plant materials and bending treatment

Experiments were carried out on hybrid poplars (*P. tremula* × *P. alba*, clone INRA 717-1B4). Poplar plants were regenerated through *in vitro* micropropagation on MS Medium and were progressively acclimatized in hydroponic solution in the growth chamber under controlled conditions [22 °C, 60% relative air humidity, 16:8-h (L:D) photoperiod, and 70 μmol.m^−2^.s^−1^ photosynthetic active radiation]. Bending treatment was applied to the 3-month-old poplars by bending of the basal stem part transiently (6 s) against a plastic tube as described by Martin *et al*. (2010). 1.5 % uniform maximal deformation was applied to all plants using the following formula: *ε*max = *r*/(*r* + *ρ*), where *ε*max is the maximal strain applied to the stems, *r* is the radius of the stem in the direction of bending, and *ρ* is the radius of the plastic tube (Roignant *et al*., 2018). For western blot and qRT-PCR analyses, stem samples were directly harvested in liquid N2 at various time points after bending treatment. For ChIP assay, stem tissues were crosslinked in formaldehyde prior to harvest in liquid N2.

### Nuclear protein extraction

Nuclear proteins were extracted from frozen poplar stem using the CellLytic™ PN kit (Sigma, USA) according to the manufacturer’s instruction. Briefly, 2 g of grind tissue was resuspended in 6 ml of 1X NIB buffer containing 1 mM DTT. The homogenized extract was filtered through filter mesh to remove cell debris and then centrifuged for 10 minutes at 1,260 x *g*. The pellet was resuspended in 1 ml NIBA buffer containing 1:100 (v/v) protease inhibitor cocktail (Sigma, P9599). Triton X-100 (1% final concentration) was added in the NIBA buffer to lyse cell membrane and then the lysate was centrifuged at 12,000 x *g* for 10 min at 4 °C. The supernatant containing the cytosolic proteins was stored at -80 °C and the nuclei pellet was washed again by resuspending in 1 ml NIBA buffer. After centrifugation, the nuclei pellet was resuspended in freshly prepared extraction buffer (2/3 of the pellet volume) containing 5 mM DTT and 1:100 (v/v) protease inhibitor cocktail. After vortexing for 30 min and spinning at 12,000 x *g* for 10 min at 4 °C, the supernatant (i.e., nuclear proteins) was collected into a new tube and stored at -80 °C. Concentration of the nuclear and cytosolic proteins were determined with the Pierce™ BCA protein assay kit (ThermoFisher, USA). Pre-cooled buffers and equipment were used for the experiment, and all steps were carried out at 2–8 °C.

### Western blotting

Equal amounts (10 µg) of nuclear proteins from different samples were separated using SDS– PAGE and electrotransferred to a nitrocellulose membrane (BioRad, 162-0115) for 1.5 h at constant 400 mA at 4 °C. The membrane was immersed in TBST solution [20 mM TRIS, 137 mM NaCl, and 0.1% Tween-20] containing 5% skimmed milk for 1h at room temperature for blocking. Next, the membrane was incubated with primary antibody (diluted in TBST solution) at room temperature for overnight. Then the membrane was rinsed with TBST solution for three times (10 min each) and incubated with a horseradish peroxidase-conjugated secondary antibody (diluted in TBST solution containing 5% skimmed milk) for 1 h at room temperature. After brief washing (3 times with TBST solution and 1 time with TBS), the blot was developed using Immobilon Western Chemiluminescent HRP substrate (Millipore, WBKLS0500), and the signal was exposed with X-ray film. Six different commercial primary antibodies were used: anti-H3pan (Diagenode, C15200011), anti-H3K9/14ac (Diagenode, C15410005), anti-H3K4me3 (Diagenode, C15410030), anti-H3K27me3 (Diagenode, C15410195; Millipore, 07-449), and anti-H3K36me3 (Diagenode, C15410192). The reactivity and specificity of these antibodies were tested using nuclear and cytosolic extracts of poplar proteins, and we found anti-H3pan, anti-H3K9/14ac, and anti-H3K4me3 antibodies are specific to common epitopes of poplar H3 histones, while others are non-specific (data not shown). The secondary antibody used against anti-H3Pan was a rabbit anti-mouse IgG (Sigma, A9044) and the secondary antibody used against other primary antibodies was a goat anti-rabbit IgG (4052-05, SouthernBiotech).

### Chromatin immunoprecipitation (ChIP)-quantitative real-time PCR

The Diagenode’s Universal Plant ChIP-seq kit (Cat. No. C01010152) was used as per the manufacturer’s instructions for a rapid and efficient ChIP analysis. The bent portion (~4 cm, 0.3-0.5 g) of the poplar stem was debarked, and then cross-linked with 1% formaldehyde.

Samples were vacuum infiltrated for 25 min (5 min x 5 times) at 4°C. The cross-linked chromatin was sheared using Bioruptor sonication system (Diagenode) and sheared chromatin was immunoprecipitated using 1 μg of antibody per reaction. For immunoprecipitation, following antibodies were used: anti-H3pan (Diagenode, C15200011), anti-H3K9/14ac (Diagenode, C15410005), anti-H3K4me3 (Diagenode, C15410030), and IgG (Diagenode, C15410206). Three biological replicates were used for each ChIP reaction. The enrichment profiles of histone modifications at regulatory regions of five genes were tested by SYBR Green–based qPCR. Primer details are available in **Table S1** and distribution of primers at regulatory regions are shown in **Figure S1** (available as Supplementary data). The qPCR reaction was carried out using 1:20 diluted immune-precipitated DNA (IP) or input. ChIP qPCR data were analyzed using the 2^(-ΔΔCt)^ fold difference method (Mukhopadhyay *et al*., 2008). First, Δ*C*t value for each sample was calculated as follows: [*C*t (IP sample) − *C*t (input)]. Next, ΔΔ*C*t was calculated as follows: [Δ*C*t (bent sample) −Δ*C*t (control)]. Finally, fold difference [2^(-ΔΔCt)^] values were normalized to a reference gene (*PtaUBQ*) and H3pan and plotted in a graph.

### Quantitative real-time PCR

Total RNA from bent portion of the poplar stems was extracted using CTAB extraction buffer as described by (Chang *et al*., 1993). RNA samples were treated with RNase-free RQ1 DNase (Promega, Charbonnières-les-Bains, France) for removal of DNA contamination. RNA was quantified through NanoDrop™ (ThermoFisher Scientific, USA). cDNA was prepared with 1 μg RNA using oligo dT and SuperScript III reverse transcriptase (Invitrogen, Cergy-Pontoise, France) as per manufacturer’s instruction. qRT-PCR was performed using a StepOnePlus™ real-time PCR system and SYBR Green fluorescent dye as previously described by (Roignant *et al*., 2018). Each PCR reaction (15 μL) contained: cDNA (3 μL of a 1:30 diluted cDNA), primers (5 μm each), and 1X Takyon Rox SYBR® MasterMix dTTp Blue (Eurogentec, Angers, France). Primers details are available in **Table S2** (available as Supplementary data). Specificity of primers for each gene was checked by analyzing individual dissociation/melting curve and running the amplicons in agarose gel. The Quantitative Relative Normalized (QRN) abundance of each transcript was calculated by comparing bent vs control plants as described by (Pfaffl, 2001). *PtaUBQ* and *PtaTIP41* were used as reference genes for normalization. In addition, *PtaUP2* and *PtaEF1* were also used as reference genes in some cases. Three biological replicates were used to calculate mean values.

### Histone deacetylase (HDAC) inhibitor treatments

Poplar plants were produced and grown under the similar conditions as mentioned above except that each plant was grown in individual plastic container which contains 5 L hydroponic solutions. HDAC inhibitor experiments were performed with 3-month-old poplars that were treated with 1.3 μM of trichostatin A (TSA, dissolved in DMSO) or only DMSO (0.07% v/v) for 6 days. After adding TSA or DMSO in hydroponic solution, the volume of solution (5 L) in each container was maintained throughout the experiment by adding extra hydroponic solution in every 2 days intervals. After 6 days, bending treatment was applied to plants and stem samples were harvested in liquid N2 at various time points. Three biological replicates were used to calculate mean values.

## RESULTS

### Global analysis of post-translational modifications of histones after bending treatment

Histone modifications within the promoter and gene body plays a key role in gene transcription. To investigate the role of histone modifications in the regulation of gene expression during the accommodation process in poplar, first, we tested the specificity of several types of commercially available antibodies against different histone methylation or acetylation marks (see method section). Only two of these antibodies, anti-H3K9/14ac and H3K4me3, showed specificity to common epitopes of poplar H3 histones, recognizing a unique protein of approximately 17kDa as expected. Nuclear proteins were extracted at various time points (1, 5, 24, and 72h) after bending treatment, and western blotting was performed to follow the level of total H3K9/14ac and H3K4me3. As shown in **Figure 1**, whereas no major changes could be detected for the H3pan and H3K4me3, we observed a reduction in the global amount of H3K9/14ac in poplar stems 24 h after bending treatment compared to control. The proportion of H3K9/14ac returned to the higher level at 72 h post bending. Alteration in the level of such N-terminal mark of histone H3 in the mechano-stimulated plants may contribute to the accommodation process.

**FIGURE 1:**
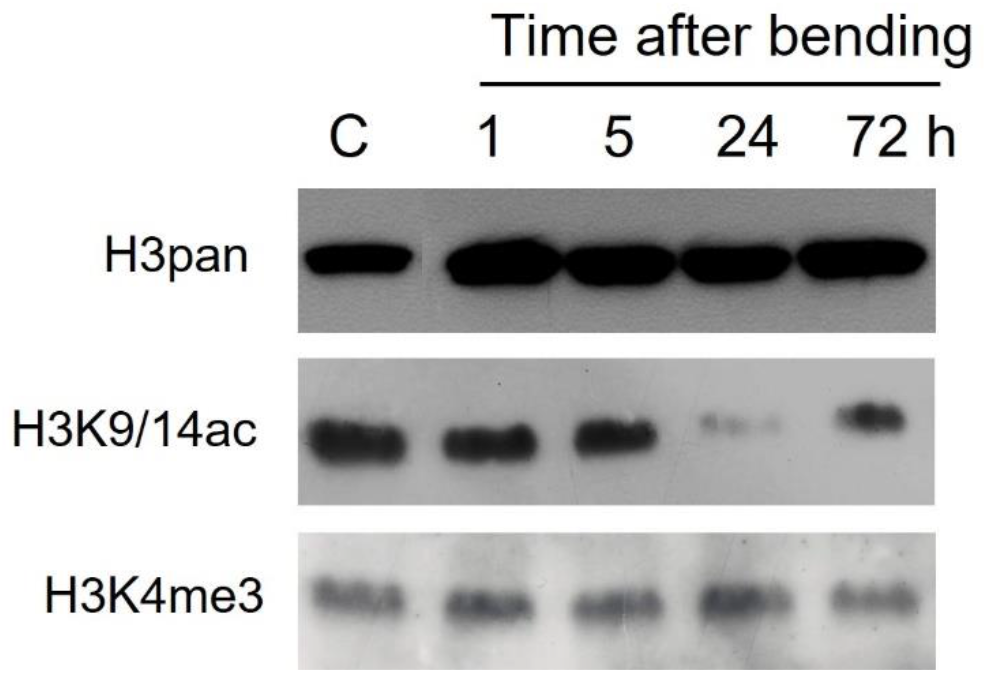
Western blot analysis of nuclear protein extracts from stems of bent and control (C) poplar plants using antibodies directed to H3pan, H3K9/14ac, and H3K4me3. The time (1, 5, 24, and 72 h) indicates h post single transitory bending of the stem. Nuclear proteins were extracted from the bent region of the stem. The same membrane was used for the hybridization with anti-H3K9/14ac followed by anti-H3K4me3, and the membrane was de-hybridized before treating with anti-H3K4me3.

### Single stem bending alters the expression pattern of genes involved in epigenetic modifications in poplar

Next, we re-analyzed the transcriptomic data available in Pomies *et al*. (2017) to find if the expression patterns of any epigenetics-related genes correlate with the kinetics of the desensitization process in the mechano-stimulated poplar. As shown in the heat map (see **Figure S2** available as Supplementary data), six genes potentially involved in the histone post-translational modifications were found to be differentially expressed in the mechano-stimulated poplar. Among them, 3 genes are associated with polycomb repressive complex (PRC): *Potri*.*018G076500* (which encodes VIN3-LIKE PROTEIN 1, VIL1), *Potri*.*014G120100* (encodes a SET domain containing histone-lysine N-methyltransferase, EZH2), and *Potri*.*019G044400* (encodes a chromo domain-containing protein, LHP1). Other 3 genes are *Potri*.*005G195400* (encodes a SET domain containing histone-lysine N-methyltransferase), SDG26 *Potri*.*009G149400* (encodes a histone deacetylase 2A, HD2A), and *Potri*.*003G096100* (encodes a JMJ family protein that acts as a lysine-specific demethylase, JMJ). Moreover, a structurally divergent linker histone H1.3 (*Potri*.*007G014200*) was noted to be differentially expressed in the bent stem. Some of these genes not only were significantly up-regulated after one bending but also showed an attenuated response after the second bending applied 24h interval (**Figure S2**). To understand better the temporal expression kinetics of these genes within the first 24h of single bending, when plants enter the desensitization phase, we performed a qPCR analysis with samples from bent stems that were harvested at 1, 5, 9, 13, 17, and 21 h after a single bending treatment (**Figure 2**). Among them, *PtaJMJ* and *PtaVIL1* were transiently up-regulated at 1 h and then returned to the basal level at 5 h after treatment. Transcript levels of *PtaHD2A* and *PtaH1*.*3* peaked at later time points, at 5 h and 9 h, respectively. The other three genes (*PtaLHP1, PtaEZH2*, and *PtaSDG26*) were mainly downregulated after bending treatment. Broadly, these results revealed that a single transitory bending of a stem regulates the expression of several genes involved in histone posttranslational modification. Such regulation could alter the chromatin accessibility of mechano-responsive genes before the second bending, followed by triggering an attenuated response.

**FIGURE 2:**
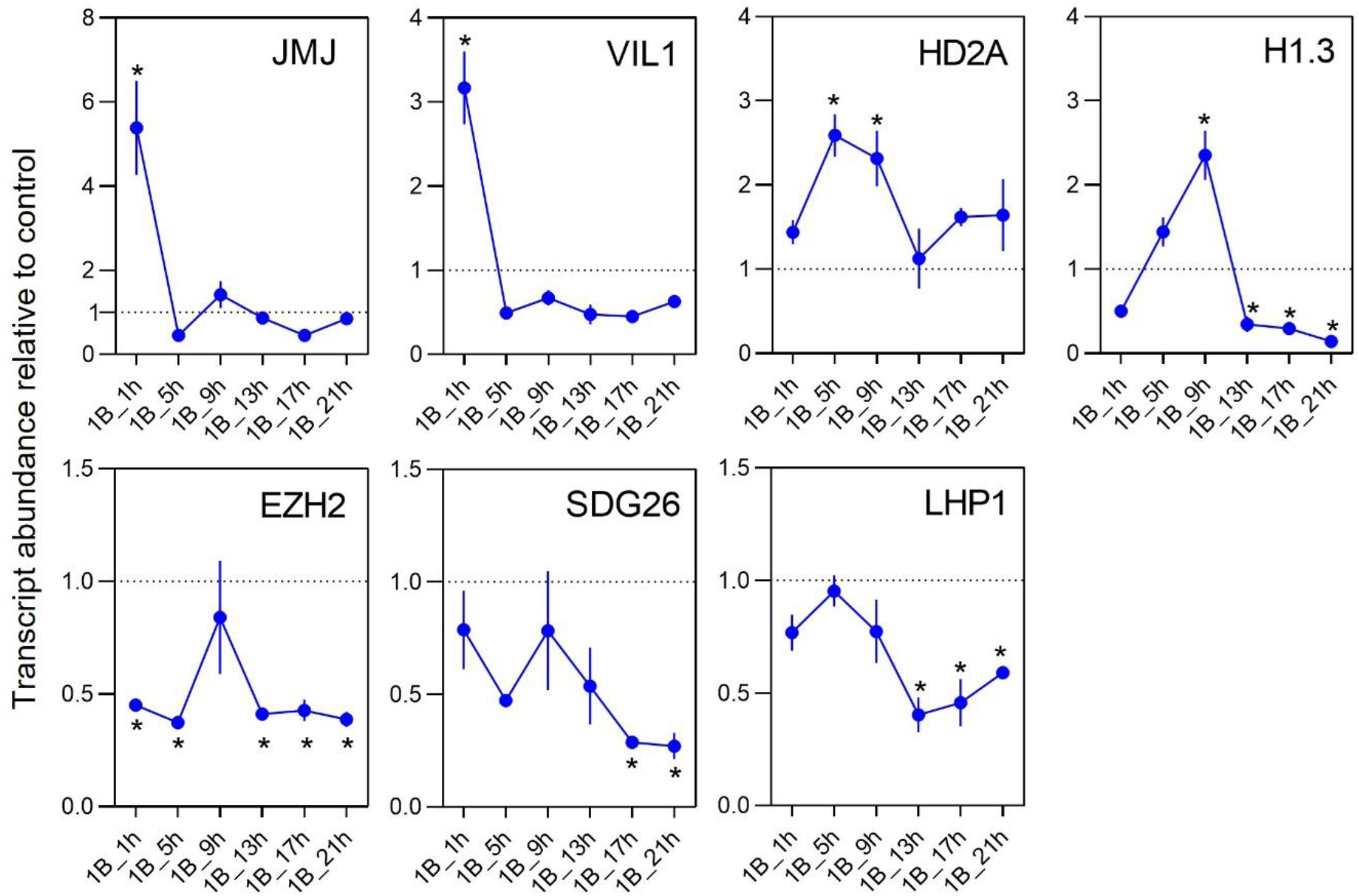
Relative transcript abundance of *PtaJMJ, PtaVIL1, PtaHD2A, PtaH1*.*3, PtaEZH2, PtaSDG26, PtaLHP1* genes after a single transient bending. The *y* axis denotes transcripts abundance relative to control (no bent, dotted line) and it was calculated after normalization using 4 reference genes (*UBQ, TIP41, UP2*, and *EF1*) by qPCR. The time (1, 5, 9, 13, 17, and 21 h) indicates h post stem bending. Error bar indicates the standard error of means (*n*=4). The asterisks (*) represent a significant difference compared to the control at the significance level of 5% (Kruskal and Wallis test).

### Comparison of expression patterns of two types of mechano-responsive genes during desensitization and resensitization phases

Transcriptome analysis by Pomies *et al*. (2017) revealed two types of transcriptional regulation of mechano-responsive genes by repetitive stimulation with a 24h interval: genes that are identically regulated after one or two bending (non-attenuated genes) and genes which show attenuated expression during the second bending (attenuated genes). To compare the kinetics of histone modifications in the regulatory region of these two types of genes during the desensitization-resensitization processes, we first chose four genes from the microarray data to analyze their expression kinetics more precisely, especially during the resensitization phase that was not analyzed during the microarray analysis. Three of them were chosen for their typical attenuated expression pattern (*PtaZFP2, PtaXET6*, and *PtaACA13*), and the fourth one, *PtaHRD*, is a non-attenuated gene (see **Figure S2** available as Supplementary data). We also checked the expression pattern of two potential histone modifiers, *PtaJMJ* and *PtaVIL1*, which were noted to be transiently up-regulated in the bent stem (**Fig. 2)**. The resensitization process was analyzed by applying two transitory bending treatments separated by long intervals, such as 3 or 5 days (see **Figure S3A** for sample harvest strategy). To do so, RNA from the mechano-stimulated poplar stems were isolated at the following time points: 30 min after 1 bending (1B_30m), 3 days after 1 bending (1B_3d), 5 days after 1 bending (1B_5d), second bending applied with a 3-days interval (2B-3di), and second bending applied with a 5-days interval (2B-5di). Twice mechanostimulated samples (i.e., 2B-3di and 2B-5di) were harvested 30 min after the second bending. The control sample was harvested at the beginning as well as end (i.e., on the 5th day) of the experiment, and transcript levels in the bent zone of stem were quantified relative to the average of the two controls. qPCR analysis shows that all 4 genes were highly up-regulated 30 min after a single bending and returned to a basal level after 3 and 5 days (**Figure 3**). When second bending was applied 3 days after the first one, the expression of *PtaZFP2, PtaXET6, PtaACA13, PtaJMJ*, and *PtaVIL1* was markedly weaker. However, when the second bending was applied with a 5-days interval, these five genes were strongly up-regulated similarly to that of 1B_30m sample. In contrast, *PtaHRD* did not show any attenuated expression pattern after the second bending applied with a 3-days interval. Moreover, our result showed that the *PtaHRD* upregulation was even stronger in 2B-3di and 2B-5di samples compared to the control. Collectively, these results validate the kinetics of the accommodation process in poplar, that the desensitization phase, which occurs within 24h of mechanostimulation, lasts for several days.

**FIGURE 3:**
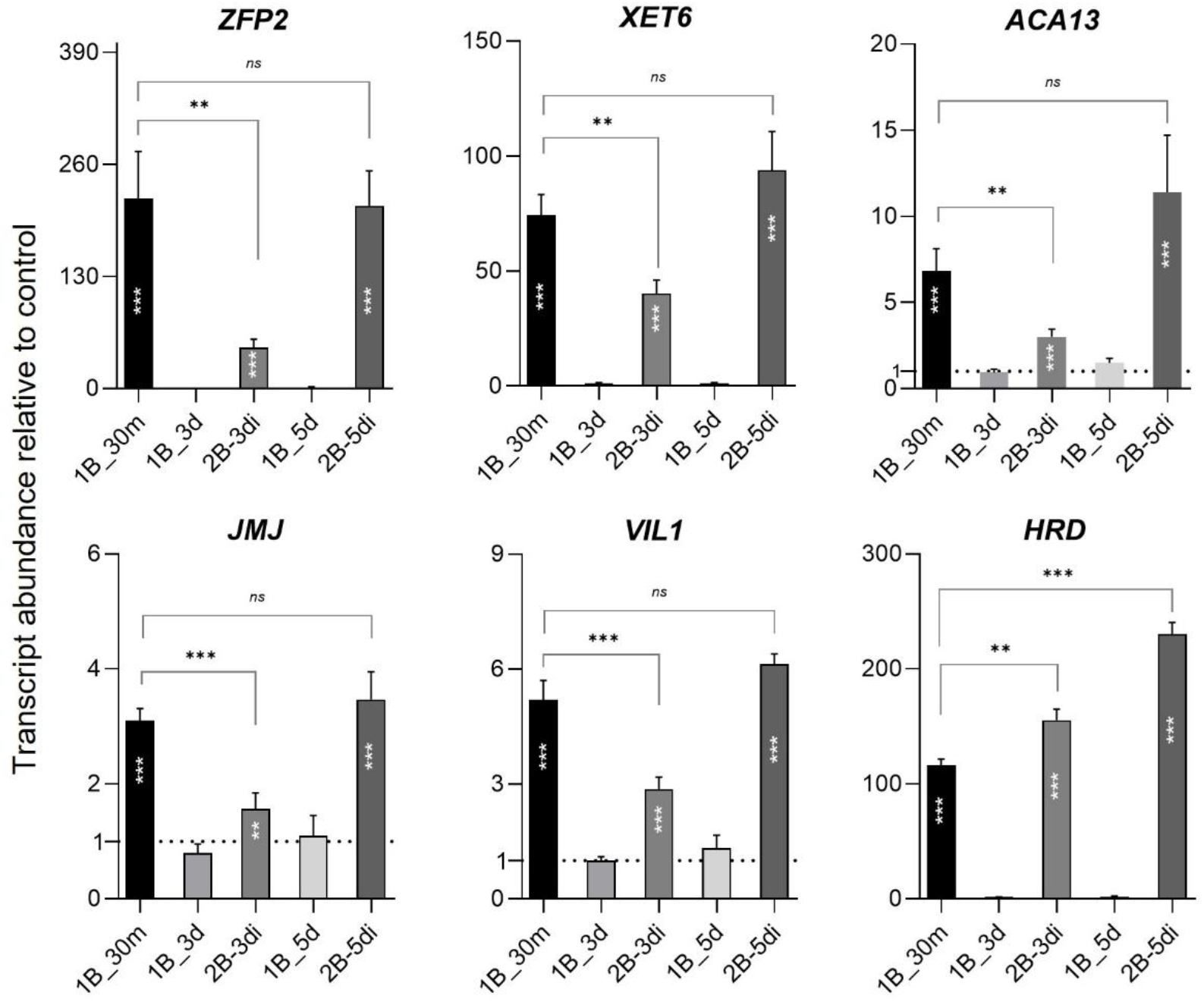
Relative transcript abundance of *PtaZFP2, PtaXET6, PtaACA13, PtaHRD, PtaJMJ*, and *PtaVIL1* genes after repeated stem bending. Two bending treatments were applied at 3-or 5-days intervals. Total RNAs were extracted from stems of control (no bent), 1B_30m (harvested 30 min after 1 bending), 1B_3d (harvested 3 d after 1 bending), 1B_5d (harvested 5 d after 1 bending), 2B-3di (harvested after 2 bendings applied with a 3-d interval), and 2B-5di (harvested after 2 bendings applied with a 5-d interval). The *y* axis denotes transcripts abundance relative to control (set to 1, dotted line) and it was calculated after normalization using 2 reference genes (*UBQ* and *TIP41*) by qPCR. Error bar indicates the standard error of means (*n*=3). Statistical significance was determined by Student’s T-test and *P*-value ranges are marked by asterisks: *** *P* < 0.01, ** 0.01 < *P* < 0.05, * *P* < 0.1. ns: non-significant. Asterix within bar indicates the significant differences with control.

### The levels of H3K9/14ac in the regulatory region of attenuated genes correlate well with the desensitization phenomenon

The H3K9/14ac mark is typically associated with the actively transcribed genes in their proximal promoters and exon (Singh *et al*., 2014). In order to check whether the level of H3K9/14ac in the mechanoresponsive loci correlates with their expression pattern during the desensitization and resensitization phases, we analyzed the level of H3K9/14ac mark in the attenuated (*PtaZFP2, PtaXET6, PtaACA13*) as well as non-attenuated (*PtaHRD*) loci using ChIP assay. Broadly, ~1 kb nucleotide sequence around the translation start site (ATG) of these genes were extracted from the URGI database (i.e., whole-genome draft assembly of *P. tremula* x *P. alba* clone INRA 717-1B4). The sequences of these genes were further confirmed by sanger sequencing (see **Table S3** available as Supplementary data). For CHIP qPCR analysis, three sets of primers per gene were designed targeting the following regions (**Figure S1**): distal (P1) and proximal (P2) regions of the upstream of the start codon, and the first exon (P3). Next, we measured the level of H3K9/14ac after applying two transitory bending treatments with 3-or 5-days intervals. Like the qPCR analysis as shown in **Figure 3**, samples for the ChIP assays were harvested at seven different time points (see **Figure S3A** for sample harvest strategy). As shown in **Figure 4**, the strongest enrichment of H3K9/14ac marks was observed at 30 min after the first bending in the following regulatory regions of attenuated genes: P1 and P2 of *PtaZFP2*, P2 of *PtaXET6*, and P2 and P3 of *PtaACA13*. However, 3 and 5 d after the first bending, the level of H3K9/14ac marks was reduced and returned to the basal level in those regions. As expected with the expression patterns, in comparison to 3 days after the first bending (1B_3d), the level of H3K9/14ac was not significantly increased after the second bending applied with a 3-days interval (2B-3di) during the desensitization phase. When second bending was applied with a 5-days interval (2B-5di) during the resensitization phase, we observed 1B_30m-like enrichment in the P1 region of *PtaZFP2* and P2 region of *PtaACA13* but not in the P2 region of *PtaZFP2* and *PtaXET6* and the P3 region of *PtaACA13*. Broadly, the H3K9/14ac status in the regulatory regions of attenuated genes correlates well with their expression pattern after the second bending at the desensitization phase but not at the resensitization phase. It suggests that the regulation of H3K9/14ac marks is not sufficient to explain the resensitized expression of these three attenuated genes. Nevertheless, together with the western blotting result, the ChIP assay indicates the critical role of histone acetylation in the expression of attenuated genes after mechanostimulation. In contrast, the enrichment level of H3K9/14ac marks was not strongly different in the *PtaHRD* loci after 1 or 2 successive bending, irrespective of the interval between two treatments (**Figure 4**), which could differentiate it as a separate class of regulation from the attenuated genes.

**FIGURE 4:**
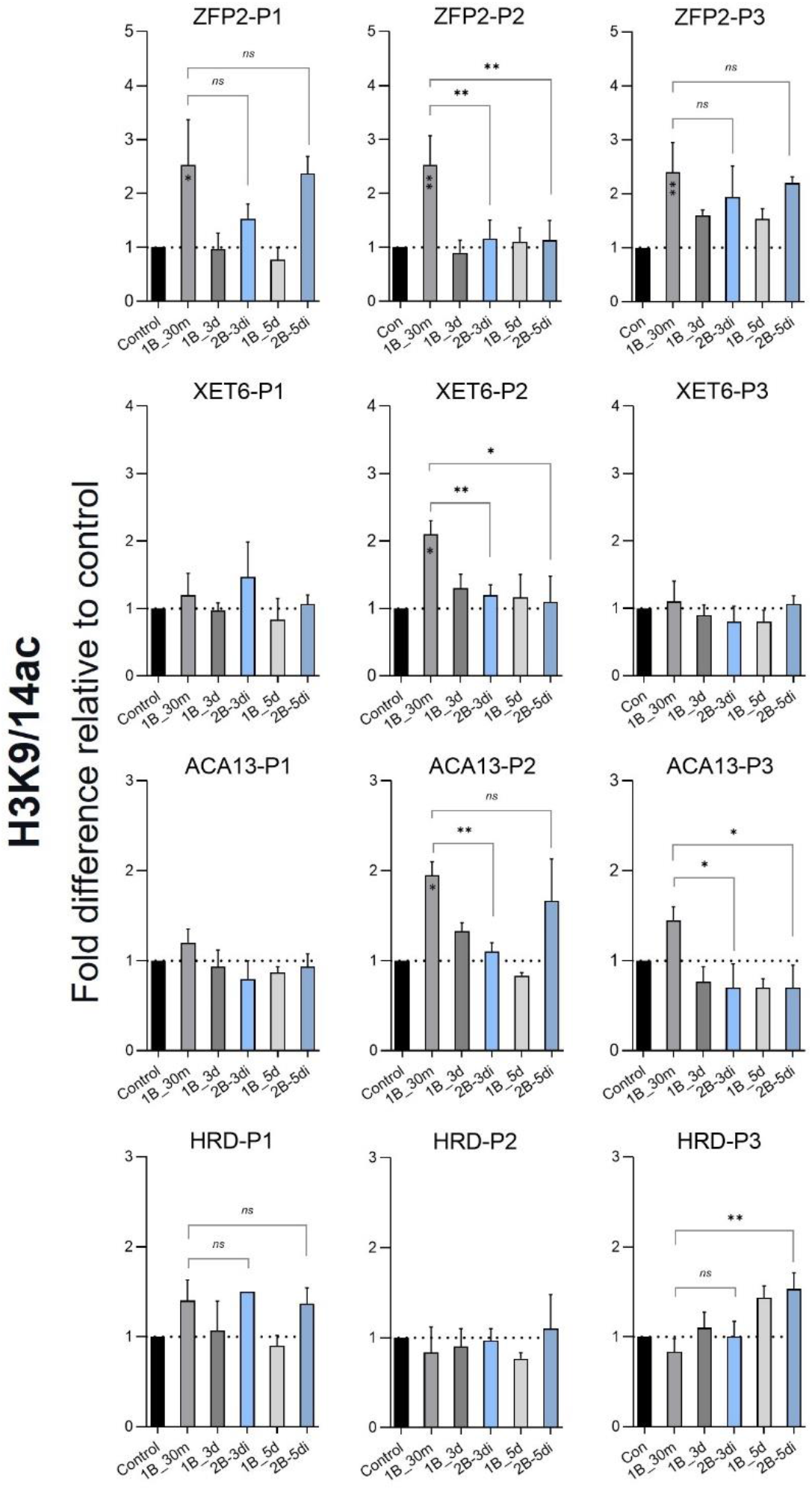
Dynamics of H3K9/14ac modification at *PtaZFP2, PtaXET6, PtaACA13*, and *PtaHRD* after applying two bending treatments at 3-or 5-days intervals. Chromatin was extracted from stems of control (no bend), 1B_30m (harvested 30 min after 1 bending), 1B_3d (harvested 3 d after 1 bending), 1B_5d (harvested 5 d after 1 bending), 2B-3di (harvested after 2 bendings applied with a 3-d interval), and 2B-5di (harvested after 2 bendings applied with a 5-d interval) plants. ChIP was performed with antibodies against H3K9/14ac and H3pan. Primers for qPCR were designed targeting the following regions (see **Figure S1** for schematic representation): distal (P1) and proximal (P2) regions of the upstream of the start codon, and the first exon (P3). Amplification values were normalized to input. The *y* axis denotes fold difference relative to control (set to 1, dotted line) and it was normalized to H3pan and P2 region of *UBQ*. Error bar indicates the standard error of means (*n*=3). Asterix within bar indicates the significant differences with control determined by one-way ANOVA with Dunnett’s multiple comparisons test. Statistical significance between 1B_30m and 2B-3di or 2B-5di was determined by Student’s T-test. *P*-value ranges are marked by asterisks: *** *P* < 0.01, ** 0.01 < *P* < 0.05, * *P* < 0.1. *ns*: non-significant.

### Expression efficiency of the attenuated genes was restored after second bending in the TSA-treated plants

We wanted to confirm whether the global reduction of H3K9/14ac mark at 24 h post bending, as observed in **Figure 1**, is responsible for the attenuated expression pattern after the second bending at the desensitization phase. Therefore, we compared the expression pattern of selected attenuated and non-attenuated genes in the normal and TSA-treated plants after repetitive bending treatment with a 24 h interval. First, the desensitization process in normal plants was confirmed by quantifying the transcripts level of eight mechano-responsive genes. For this purpose, one or two transitory stem bending was applied with a 24 h interval, and stem samples were harvested 30 min after treatment (see **Figure S3B** for sample harvest strategy). The control sample (i.e., without bending) was harvested at the end of the experiment, and transcript levels of genes in the bent zone of the stem were quantified relative to the control samples. The results confirm that transcript accumulation of *PtaZFP2, PtaXET6*, and *PtaACA13* were markedly weaker after the second bending relative to the first bending (**Figure 5A**). To confirm this accommodation phenomenon, we measured the transcript levels of two important mechano-signalling genes. Among them, Feronia (FER) receptor-like kinase mediates the mechanotransduction process in Arabidopsis (Shih *et al*., 2014). Several *calmodulin-like* (*CML*) genes, including touch marker genes *TCH2* and *3*, were noted to be upregulated in the touch-treated Arabidopsis (Lee *et al*., 2005). Both the *PtaFER* and *PtaCML11* showed attenuated expression patterns after the second bending. We also checked the expression pattern of two potential histone modifiers, *PtaJMJ* and *PtaVIL1*. The expression of these two genes was attenuated when two bendings were separated with a 24-h interval (**Figure 5A**). We could not test it for the other five epigenetics-related genes as they were poorly expressed at 30 min after 1 bending (see **Figure S4** available as Supplementary data). On the other hand, as expected, the upregulation pattern of *PtaHRD* remained unaltered after twice bending treatment relative to the first one.

**FIGURE 5:**
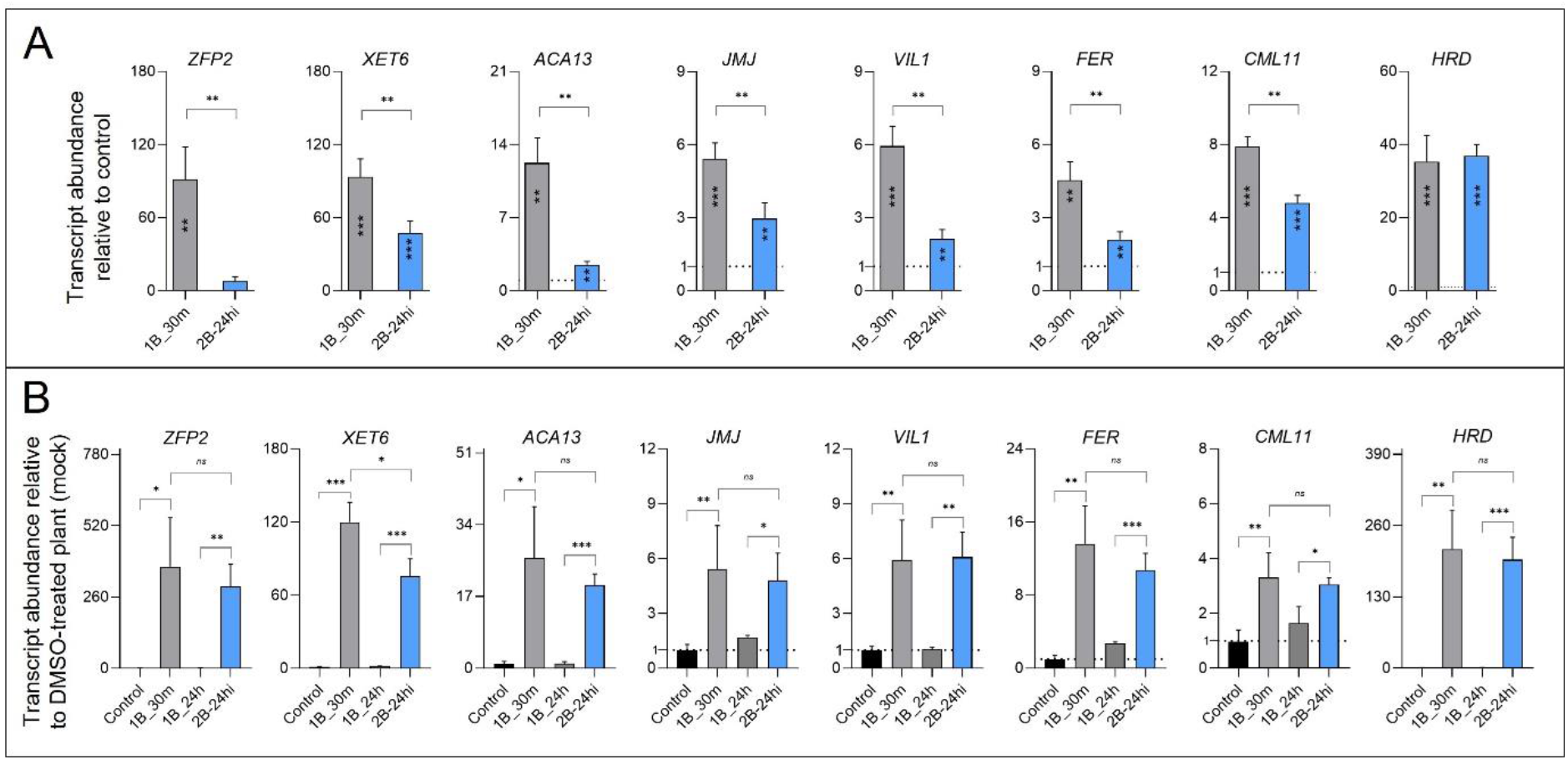
Relative transcript abundance of attenuated and non-attenuated genes in the normal and TSA-treated plants after repeated stem bending. **A)** Poplar plants were subjected to one or two stem bending treatments with a 24 h interval. The *y* axis denotes transcripts abundance relative to control (set to 1, dotted line). Asterix within bar indicates the significant differences with control. **B)** Trichostatin A-treated plants were subjected to one or two bending treatments with a 24 h interval. The *y* axis denotes transcripts abundance relative to DMSO-treated plant (mock, set to 1). Total RNAs were extracted **from** stems of control (no bent), 1B_30m (harvested 30 min after 1 bending), 1B_24h (harvested 24h after 1 bending), and 2B-24hi (harvested after 2 bendings applied at 24 h intervals) plants. Reference genes *UBQ* and *TIP41* were used for normalizing the qPCR data. Error bar indicates the standard error of means (*n*=3). Statistical significance was determined by Student’s T-test and *P*-value ranges are marked by asterisks: *** *P* < 0.01, ** 0.01 < *P* < 0.05, * *P* < 0.1. ns: non-significant.

Next, we checked the expression pattern of these genes in the mechanostimulated stem treated with TSA. TSA inhibits the activity of histone deacetylase (HDAC), an enzyme that removes acetyl groups from lysine of both histone and non-histone proteins (Zhang *et al*., 2013). For gene expression analysis, samples were harvested as follows: control plants (TSA-treated, no-bend), 1B_30m (TSA-treated, harvested 30 min after 1 bending), 1B_24h (TSA-treated, harvested 24 h after 1 bending), and 2B-24hi (TSA-treated, harvested 30 min after second bending applied at 24 h intervals). Transcript levels in TSA-treated samples were quantified relative to the DMSO-treated plants (mock). In the presence of TSA, no significant difference was observed in the upregulation pattern of attenuated genes *PtaZFP2, PtaACA13, PtaJMJ, PtaVIL1, PtaFER* and *PtaCML11* after once or twice bending treatment (**Figure 5B**), mimicking the non-attenuated *PtaHRD* gene expression pattern. This result indicates that inhibiting histone deacetylase reduces the desensitization phenomenon, and that lysine acetylation status might play a significant role in the mechano-stimulated expression of attenuated genes.

### Levels of H3K4me3 in the regulatory region of attenuated genes correlate well with the de- and re-sensitization phenomenon

H3K4me3 is generally associated with the 5’ end of coding region of actively transcribed genes (Kim *et al*., 2008; Zhang *et al*., 2009). Thus, we performed ChIP-qPCR only with the primers specific to the P3 region of the different loci. Like H3K9/14ac, enrichment of H3K4me3 was noted at 30 min after the first bending in all three attenuated loci, and later it was reduced to basal level 3 or 5 days after the first bending (1B_3d and 1B_5d samples) (**Figure 6**). As expected, when applying the second bending during the desensitization phase (2B-3di), the level of H3K4me3 was significantly less relative to the first bending (1B_30m). In fact, no enrichment of H3K4me3 marks was observed at 2B-3di in comparison to 1B_3d. However, when the second bending was applied during the resensitization phase (2B-5di), the mark was strongly increased relative to 1B_5d. Also, we observed 1B_30m-like or higher enrichment of H3K4me3 when second bending was applied with a 5-days interval. Broadly, the enrichment level of H3K4me3 correlates well with the expression pattern of the attenuated genes during the desensitization as well as resensitization phase. Again, the pattern of H3K4me3 modifications in the non-attenuated *PtaHRD* locus differs from those of attenuated genes. It correlates well with the expression pattern of *PtaHRD* since the level of the H3K4me3 mark increased between one and two bending (**Figure 6**).

**FIGURE 6:**
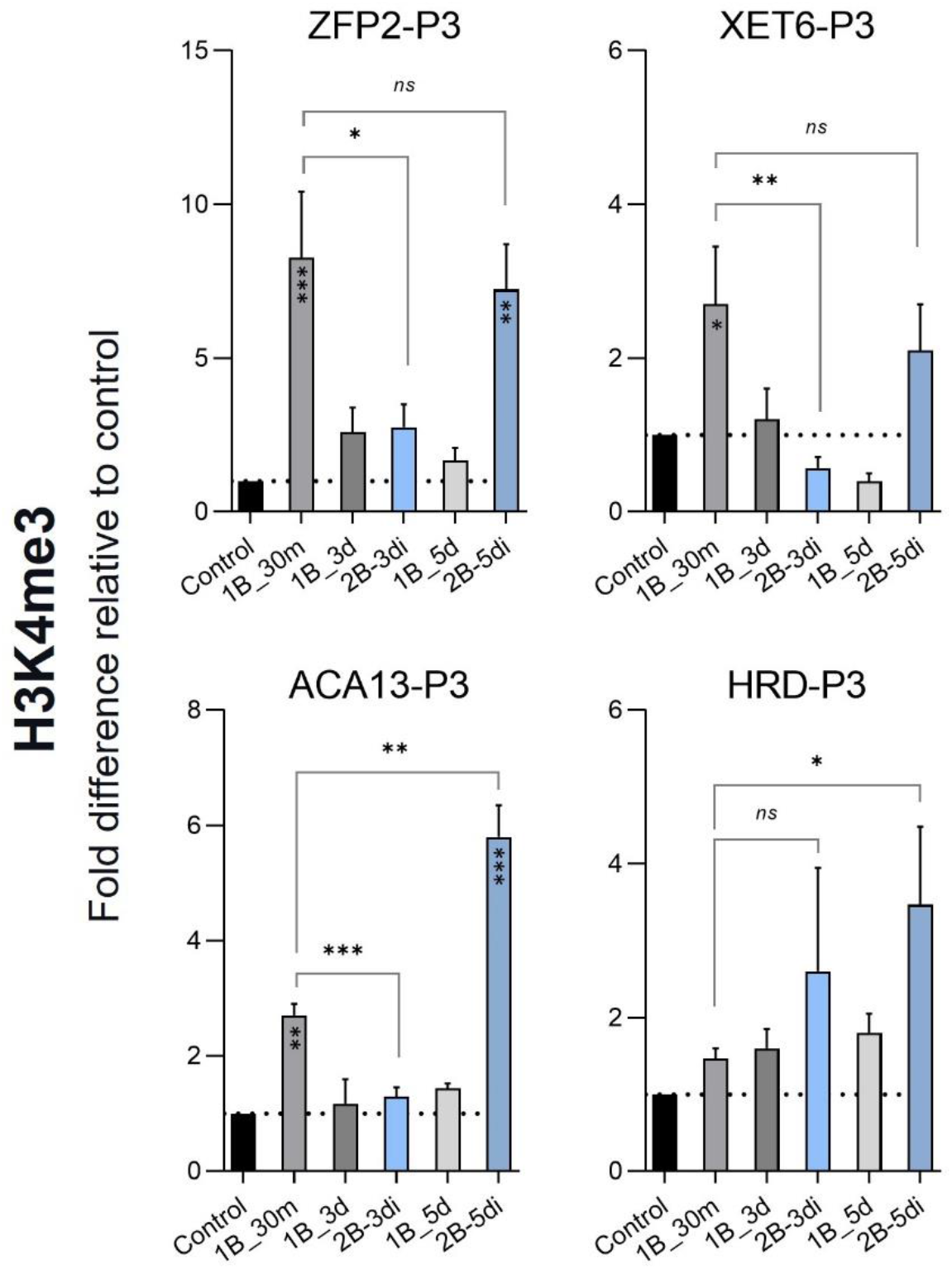
Dynamics of H3K4me3 modifications at *PtaZFP2, PtaXET6, PtaACA13*, and *PtaHRD* after applying two bending treatments at 3- or 5-days intervals. Chromatin was extracted from stems of control (no bent), 1B_30m (harvested 30 min after 1 bending), 1B_3d (harvested 3 d after 1 bending), 1B_5d (harvested 5 d after 1 bending), 2B-3di (harvested after 2 bendings applied with a 3-d interval), and 2B-5di (harvested after 2 bendings applied with a 5-d interval) plants. ChIP was performed with antibodies against H3K4me3 and H3pan. qPCR was performed only with the primers specific to the first exon P3 region (see **Figure S1** for schematic representation). Amplification values were normalized to input. The *y* axis denotes fold difference relative to control (set to 1, dotted line) and it was normalized to H3pan and P2 region of *UBQ*. Error bar indicates the standard error of means (*n*=3). Asterix within bar indicates the significant differences with control determined by one-way ANOVA with Dunnett’s multiple comparisons test. Statistical significance between 1B_30m and 2B-3di or 2B-5di was determined by Student’s T-test. *P*-value ranges are marked by asterisks: *** *P* < 0.01, ** 0.01 < *P* < 0.05, * *P* < 0.1. ns: non-significant.

## DISCUSSION

Trees are exposed to a fluctuant mechanical environment throughout their perennial life, which they can cope with by acclimating their growth. The current study confirms the accommodation of transcriptional response in poplar during repetitive mechanostimulation, highlighting the fact that plants can fine-tune their responses to mechanical stimulation. In poplar, it is clear that an interval of 5 days between two successive bending events is necessary to restore the expression potential of attenuated genes. Based on the outcome of this study, we can consider *PtaZFP2, PtaXET6, PtaACA13, PtaJMJ*, and *PtaVIL-1* as molecular markers of the accommodation process. Homologs of these genes could be used to understand the desensitization-resensitization phenomenon in other mechanostimulated plants. On the other hand, *PtaHRD* behaves like a typical “memory gene” that shows hyper induction upon recurrent stress relative to the first stress, as we often observe in the case of priming responses. This gene encodes an AP2/ERF-like transcription factor that has a role in increasing tolerance to salt and water stresses in both rice and Arabidopsis (Karaba *et al*., 2007). The molecular mechanisms behind such transcriptional response remain elusive in the case of mechanotransduction pathways, but H3K4me3 mark plays a vital role in memory gene expression in other abiotic stresses (Baurle and Trindade, 2020). For instance, dehydration memory genes (e.g., *RD29B* and *RAB18*) were noted to be associated with higher H3K4me3 in the repetitively dehydrated plants than plants stressed once, which leads to hyper induction upon recurrent stress (Ding *et al*., 2012). Interestingly, the enriched H3K4me3 status of these genes caused by dehydration stress can persist during the memory phase, when they produce transcripts at a basal level (Ding *et al*., 2012). Similarly, we also observed stable and moderately elevated levels of H3K4me3 at *PtaHRD*-P3 30 min, 3 days, and 5 days after one bending. Moreover, repetitively bent plants displayed stronger enrichment of H3K4me3 at *PtaHRD*-P3 than plants bent once, which correlates well with its increased transcript level after two bendings separated by 3-or 5-days intervals. On the contrary, in the case of attenuated genes, a reduced level of H3K4me3 mark could be responsible for their lower transcript abundance after two bendings with a 3-days interval. On the other hand, 1B_30m-like enrichment of H3K4me3 at attenuated loci was observed when second bending was applied 5 days after the first one. In Arabidopsis, it has been noted that SDG8, a histone lysine methyltransferase, is required for maintaining the H3K4me3 mark at *TCH3* locus and for their full mechano-responsive expression (Cazzonelli *et al*., 2014). Interestingly, we also observed reduced expression of a homologous *PtaSDG26* at 17 and 21 h post bending, which may lower the H3K4me3 level at attenuated loci when the second bending was applied with a 3-days interval.

Acetylation of histone tails is normally associated with actively transcribed genes, while deacetylation by histone deacetylase (HDAC) represses gene transcription (Kumar *et al*., 2021). Multiple studies showed that stress treatment induces H3K9/14ac levels at stress-inducible genes in plants (Chen *et al*., 2010; Singh *et al*., 2014). Similarly, we also noticed enrichment of H3K9/14ac mark immediately (30 min) after the first bending at attenuated loci, like *PtaZFP2* (P1, P2, and P3), *PtaXET6* (P2), and *PtaACA13* (P2 and P3). The same regions were found to be hypoacetylated 3 days after one bending. These results suggest that deacetylation of K9/K14 of histone H3 could be involved during the desensitization phase. Western blot analysis showed that the level of H3K9/14ac was globally reduced at 24 h of one bending and could not recover to the control level even 72 h post bending, which suggests an increasing activity of HDAC in the bent stem. Interestingly, *PtaHD2A*, a gene encoding a histone deacetylase 2A, was upregulated at 5 and 9 h post bending. An increased level of HDAC in the bent stem could affect the acetylation process even 3 days after the first bending, which in turn prevents re-enrichment of H3K9/14ac at attenuated loci after second bending at the desensitization phase. Restored expression pattern of attenuated genes in the TSA (HDAC inhibitor)-treated plants after two bendings during the desensitization phase also corroborates a prominent role of HDAC in this pattern of gene expression. Multiple studies have already established the importance of HDAC behind plant responses to environmental cues (Kumar *et al*., 2021). For instance, stress-responsive genes showed enrichment of H3K9ac at their promoters along with strong upregulation in HDAC mutants compared to wild type under salt and drought stress (Zheng *et al*., 2016). Interestingly, it has been noted that environmental stresses like wounding and exogenous application of jasmonic acid (JA) can induce HDAC transcription in *Arabidopsis* plants (Zhou *et al*., 2005). Mechanostimulation upregulates JA in *Arabidopsis* plants, and JA-signalling plays a dominant role in plant mechano-signalling (Chehab *et al*., 2012; Van Moerkercke *et al*., 2019). Therefore, detailed research with HDAC will be promising as it will broaden our understanding of plant mechano-signalling and responses. However, we could not observe 1B_30m-like enrichment of H3K9/14ac in *PtaZFP2* (P2), *PtaXET6* (P2) and *PtaACA13* (P3) regions after two bendings with a 5-days interval, although their induction pattern was restored. This result indicates that modifications of the H3K9/14ac marks cannot explain alone the desensitization-resensitization of expression pattern.

Three PRC-associated genes, *PtaVIL1, PtaEZH2*, and *PtaLHP1*, appeared to be differentially regulated in the stem bent once. Deposition of the repressive histone mark H3K27me3 by the PRC is considered one of the major gene silencing systems in plants (Schubert *et al*., 2006). The inverse deposition pattern between H3K27me3 and H3K4me3, as well as H3 acetylation, was already reported by multiple studies (Sung *et al*., 2006; Xing *et al*., 2018). Taken together, the enrichment of H3K27me3 could also contribute to the desensitization phenomenon by suppressing the expression of attenuated genes during the second bending. We could not check the status of H3K27me3 at the attenuated loci as the antibody was not specific to poplar. However, ChIP assays with H3K27me3 are warranted in future to unravel its role in accommodation process.

Together these results indicate that plants are so sensitive to mechanical cues that even 6 seconds of mild bending can enrich epigenetic marks within 30 mins of treatment, followed by rapid removal of those marks and maintaining desensitization state for several days. Based on the results obtained in our study, we propose a model depicting the role of histone modifications in the expression of attenuated genes (e.g., *PtaZFP2*) under recurrent mechanical stimulation (**Figure 7**). Enrichment of the H3K9/14ac and H3K4me3 marks associated with attenuated genes confer their upregulation immediate after the first mechanical stimulation. Later, the reduction of active marks (e.g., deacetylation of lysine by HDAC) and putative enrichment of repressive marks together attenuates their expression under recurrent mechanical stimulation in the desensitized plant. After a sufficient interval, the re-deposition of H3K4me3 (in some cases H3K9/14ac) could restore the upregulation of attenuated genes under recurrent stimulation. These results indicate the importance of a better understanding of epigenetic regulation in perennial plants that are subjected to recurrent environmental cues. In future, genetically modified poplars with altered epigenetic processes could be a promising tool to demonstrate their role behind acclimated phenotypes under recurrent environmental cues.

**FIGURE 7:**
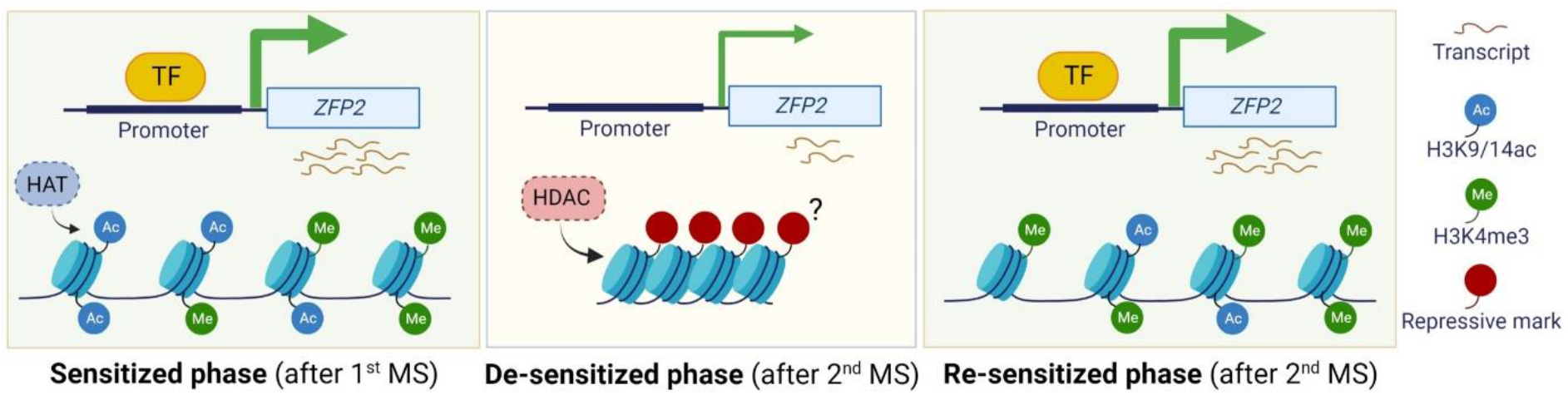
A model summarizes the role of histone modifications in attenuated gene (e.g., *PtaZFP2*) expression upon recurrent mechanical stimulation. Size of arrow (green) indicates higher or attenuated expression. Question mark indicates putative enrichment of repressive histone mark. MS: Mechanical stimulation; TF: Transcription factor; HAT: Histone acetyltransferase; HDAC: Histone deacetylase.

## Data and materials availability

The data that supports the findings of this study are available in the supplementary material of this article.

## Supplementary data

**Figure S1:** Schematics show the positions of primers for CHIP-qPCR analysis.

**Figure S2:** A heat map shows the expression kinetics of mechano-responsive genes determined by microarray analysis.

**Figure S3:** Schematic representation of experimental strategy.

**Figure S4:** Relative transcript abundance of *PtaHD2A, PtaH1*.*3, PtaEZH2, PtaSDG26*, and *PtaLHP1* genes 30 min after single transitory stem bending.

**Table S1:** List of ChIP qPCR Primers.

**Table S2:** List of qPCR primers for gene expression analysis.

**Table S3:** Nucleotide sequences that were targeted for ChIP-qPCR analysis.

## Acknowledgement

The authors are grateful to Amélie Coston for plant production and Caroline Savel for her help on RNA extraction and qPCR. Figure 7 was created with BioRender (https://biorender.com).

## Authors’ contributions

N.L.F. and R.G. conceived and planned the project. N.L.F. supervised the project. R.G., J.R., and J.F. performed experiments. R.G., N.L.F., J.R., J.F., and A.P. analyzed data. R.G. wrote original draft. R.G. and N.L.F. wrote the manuscript with input from the co-authors.

