## Supplemental Tables S1-S3 for "Temporal modification of H3K9/14ac and H3K4me3 histone marks mediates mechano-responsive gene expression during the accommodation process in poplar"

### Supplementary Tables

**Table S1**: List of ChIP-qPCR primers.

| *P. tremula x alba* | *P. trichocharpa* homologs | Primers |
|---|---|---|
| *PtaZFP2* | Potri.001G235800 | **ZFP2-F1** CACAGTGCAGTTCACAGTAAAACAT<br>**ZFP2-R1** ACTTAGAAGCGCGTGCATGATAC<br>**ZFP2-F2** TTACCAACGCTAAACTCTCACCT<br>**ZFP2-R2** GAGCAAGAACACCAAAGTATAAAGC<br>**ZFP2-F3** ATGCAAAACATGCGACAAGA<br>**ZFP2-R3** CTGGTGGTATCAACATTATTAGGC |
| *PtaXET6* | Potri.018G094900 | **XET6-F1** TTAGTCCCTGTGTTTTGAATGG<br>**XET6-R1** ACAAAACCGCCAAAACCACTAT<br>**XET6-F2** GTCAGGAATCCCATATCATTGTG<br>**XET6-R2** GCAGCAAGCTCTTGACTCTTTAC<br>**XET6-F3** GAGTCAAGAGCTTGCTGCAATAC<br>**XET6-R3** TCTGTCAAGGGAAAGTGTTAGAAG |
| *PtaACA13* | Potri.008G159100 | **ACA13-F1** CTTCTCTCTTTCTTGAGCAGGTG<br>**ACA13-R1** CTGATTTCGGGTGTTCATTATCC<br>**ACA13-F2** CTGGATAATGAACACCCGAAAT<br>**ACA13-R2** AATGCTAGAATAGACGAGGAGGAA<br>**ACA13-F3** GACGTTATGTCCCACCTTCTG<br>**ACA13-R3** TGAATCTTGAAGTGGTCAGTGTC |
| *PtaHRD* | Potri.006G021000 | **HRD-F1** GGGTTATGCTCAGTTGATACGTTA<br>**HRD-R1** TGACAGCTTTATGGTCTGAGGAT<br>**HRD-F2** CCCTAAGAAGTGATGTAAGATGTCC<br>**HRD-R2** TTCAAGTGTTCCATGAGAGAAGC<br>**HRD-F3** TTCTCTCATGGAACACTTGAAGC<br>**HRD-R3** CAGACACCCATTTTCCACTACTC |
| *PtaUBQ* | Potri.012G033000 | **UBC-F1** GAAAGAAACAGAACCATCCAATCAC<br>**UBC-R1** ATCTCTCCTCTCCTTCTGGAAATTA |

**Table S2**: List of qPCR primers for gene expression analysis.

| *P. tremula x alba* | *P. trichocharpa* homologs | Primers |
|---|---|---|
| *PtaZFP2* | Potri.001G235800 | **FP:** CGTGCGAGTCACAAGAAACC<br>**RP:** CACAGAACTCTCTTGCTGCT |
| *PtaXET6* | Potri.018G094900 | **FP**: TGTATGGGGTCTAATGGAGTC<br>**RP:** GGAATCGCTTAGTGTCTGC |
| *PtaACA13* | Potri.008G159100 | **FP:** GCCACAGTTTTGAGGTGGGGAAG<br>**RP:** AGCTAGAGCCAGAGCACCCAGTG |
| *PtaHRD* | Potri.006G021000 | **FP:** GATGCTTTAGGGATTCCAAGTATG<br>**RP:** GAATTTCCAAAGGTTCTGGTCTC |
| *PtaCML11* | Potri.008G159300 | **FP:** AGGTGGATCAGATGATTAAAGAGG<br>**RP:** CTGGCTACATTACAACAGCAACAAC |
| *PtaFER* | Potri.006G110000 | **FP:** AGGGTATTGTCGGAGTAGAC<br>**RP:** CTTGGAGTTAAGCCATCAGAG |
| *PtaJMJ* | Potri.003G096100 | **FP:** GACTCCTGCGATAACCTGGG<br>**RP:** CCTGCATTAAAGTCAGCGTTGT |
| *PtaVIL1* | Potri.018G076500 | **FP:** TGAAGAGCTGACTCCTCCATTT<br>**RP:** AGTAGGCACTTCCCCACTTG |
| *PtaHD2A* | Potri.009G149400 | **FP:** CAGATGCTAAGAAGGCCGGG<br>**RP:** GGGATAGGGAACTGACGGGAA |
| *PtaH1.3* | Potri.007G014200 | **FP:** AGGTGGAGGAGAATCCCGTT<br>**RP:** TGGACTCGACCCACTCTCAT |
| *PtaLHP1* | Potri.019G044400 | **FP:** GGGGGTCCTCATACTCAGTCA<br>**RP:** TTGTTCGCTTCTCCATTACTCTC |
| *PtaEZH2* | Potri.014G120100 | **FP:** TGAAGAGCCAAGCAGTGTTGA<br>**RP:** ACATTGTGACACTTTGCGCT |
| *PtaSDG26* | Potri.005G195400 | **FP:** AGAGCATGAGAGGGATAGCCA<br>**RP:** GCCGTGCTTGTTCAAGTGTC |
| *PtaUBQ* | Potri.012G033000 | **FP:** CCCGGCTCTAACCATATCCA<br>**RP:** GGGTCCAGCTTCTTGCAGTC |
| *PtaTIP41* | Potri.001G298500 | **FP:** TAGTGATAGTGCAAATCCTGTCA<br>**RP:** CTTACAAGTTACTGTGGACCAC |
| *PtaUP2* | Potri.002G127700 | **FP:** TATCGTCTTGTGACAATTTTTAG<br>**RP:** TCATTAGCGCCAGGACTTCC |
| *PtaEF1* | Potri.006G130900 | **FP**: GCAGATGATTTGCGTTTGC<br>**RP**: TGTAACCAACCTTCTTCAGG |

**Table S3:** Nucleotide sequences that were targeted for ChIP-qPCR analysis. Sequences were extracted from the URGI database (i.e., whole-genome draft assembly of *P. tremula* x *P. alba* clone INRA 717-1B4). Sequences highlighted in grey indicate the putative 5' UTRs, identified after pairwise alignment with *P. trichocoharpa* homologs (obtained from Phytozome V3.0 and 3.1).

| |
|---|
| ***PtaZFP2*** (Populus_717-1B4_scaffolds\|scaffold_182143 & scaffold_356255)<br>CACAGTGCAGTTCACAGTAAAACATTGATCGCTTTTAGCTATTTATTATGTATAATATATCACTTTTAATTTTTT<br>ATTGATTTTTTAAATTTACCAATTAAATATTCCTTATGCAACGTCAAGTCATTTCCAATTAACATTTATTAATAG<br>GGGTTTCTACATGAACAAATCTAAGTTGGGTATCATGCACGCGCTTCTAAGTGCGGTCGGACTTGCACTCAATAT<br>AAAATATAAGGAATTGTCAACACGTGATATGATGACCTTAGGAAGGAAGAATCGTAGTAGTGGCTCACCATCAAT<br>GACCGGATGTAATCCTGCTCACCAAGTTTGCTCAACGCGGTCACGCTAACGCATCATTACCAACGCTAAACTCTC<br>ACCTTCTATTCTACTCTTATTACATGTCTACCACACGCTCTAAAGCTCCTTAGCCCACAAGTGTTGAGGGCATAA<br>TTGCCCCTCAAGCAAAGACGTCAGTGCTTACTCGGTAGTACCAGCTCCTTCCAACCTTATAAATATATGAACTCA<br>AAGTCCCTTACTTCCATCCAACCATCAAATATCTAGCCATCTTCTTCTTTCGAAGCTTTATACTTTGGTGTTCTT<br>GCTCATCGTATAAACACAAATCATACTATATATCTAACCTGTAAGCTTAATTACCATGAAGAGAGATAGAGAACA<br>GGTAGAGGTAGACTTGGCCAAATGCTTGATGCTACTTTCTAAAGTTGGCGAAGCTGATCACGAGATACTAACTAG<br>TTATAGACCAGCAGCAGCAGCAGCAACGGCAGGGGCCGGCGCCGGGGCCGGTCGCTCATTTTCATGCAAAACATG<br>CGACAAGAATTTCCCTTCATTTCAAGCATTAGGAGGCCACCGTGCGAGTCACAAGAAACCAAAACTCATGGAATC<br>AACCGGGAACTTGTTGAAGCTGCCTAATTCGCCTTCAAAGCCAAAAACTCACCAGTGCTCCATTTGCGGCCTTGA<br>GTTTCCTCTCGGGCAAGCACTCGGAGGTCACATGAGGAGACACAGGGCGCCTAATAATGTTGATACCACCAGT |
| ***PtaXET6*** (populus_717-1B4_scaffolds\|scaffold_108083)<br>TTAGTCCCTGTGTTTTGAATGGTATTATAACTTAGTACATTATGTTTGAAAAATTATAATTTAGTCTCCACTCTT<br>GTTGATCACTATGTTGCTAAGACATTTTTTGGTTTTTTTTAATCCCAGATGTATAATTTAAAAGTATGCTGGGGT<br>TGGTGGACAACCTCGGAGGTGACTCACTTTCACCAATGAATATTGAAATGTTCATTGCGGGGCTGCTATAGTGGT<br>TTTGGCGGTTTTGTTAAATTAATGCTATAGACAAGCAACACAACTTTTTAAGCCCAGTGAAAACATAAGAAAAAA<br>ATAAAATAAGTTCTAATATTATCAAAAGAAAAGCATATGAATCATTGTTTTAAACTAGGCATGAATAATTTACAA<br>AAAAATATATGGCCTCCAGAAACTGCAGCATGTAATTGGCCTGGCTAAGTTTGTGGTCAGGAATCCCATATCATT<br>GTGCCTTGTTCATGGCATCCAGCTGTTTACTTCTCATGTCATTACACCACCCAACTTGAAGTGCTCCACAGTTGA<br>AAGGAACATTCTAATTCGCCACCATCCAGACTCCCTATAAATTGTCACAGAATTGTAACATTTTAGCAGGTCAAA<br>ATCAAACAAAACAAGTAAAGAGTCAAGAGCTTGCTGCAATACTTCATATTTGAAAACCATCAAGCTTTATTATAA<br>GTTCATGGCTGCCTGCCCTTCTGACCCAAACCCTTTACTGCTGCTGCTGATGATCTCTCTTTTGGTTGGTTCTGA<br>TTTGGTGTTGGTTGACGCTGGTTCCTTTTACCAAGATGTTGATATCATATGGGGAGACGGACGAGCTAAGATACT<br>CAACAATGGCAATCTTCTAACACTTTCCCTTGACAGAG |
| ***PtaACA13*** (populus_717-1B4_scaffolds\|scaffold_154819)<br>GCGTTACAGGAAATCCAACATAAACTCCTTATTCGAAGAACTTCCCAGAAATTTCCTAGGATTTATTTCTTCTCT<br>CTTTCTTGAGCAGGTGACCAACACAACCTATATGGTTCGGTGATGGTATTATTTTCAATCGACATAAAAGCGAAC<br>CCCGCCTCGACCTTTCCCTTCTTTCCGCGCTTTGACCGGTCAAAATACCCCAGAACTAAGATAACAATTGAACAC<br>GCTGCACCGATTCTTTGTCGTCAAGAATCCACAATACAATAATATGTACTGGATAATGAACACCCGAAATCAGAT<br>TAAGTTTTAAGATAATTAATTATAACTTCAAACTTGAGAGAAGAATCCAATCCTTGCAAGCAATCTTTTCCACCC<br>GGTTGCCACCCGTCTTTCCCACCAATACTAATTAATTGGGCCATAGCCCACGGTGCTGCTATAAAAGGAACTCGT<br>GAAACAAAGAGGGAACACCATTCTAGAATTCCTCCTCGTCTATTCTAGCATTCAAATAATACGCCTCTAGGCTTC<br>CGTTATTATTGGTTTCTTCCAAGTTCTTGAAGTTATCCTTTGCTTTGTGAAATATCATTTTAGTTTCTCTATATT<br>GTCATTTCATTTTCTGCTAGTTTTAGAGTTTCGACGTTATGTCCCACCTTCTGCATGCAAACTTGGTTTGCATTG<br>AGCGTTTACTTGACGTCCCTGCCACCCTTAGCAAACCCAACAAAAGATGGCATTTAGTTTTCGCAACTATCTATT<br>GTTCCAGGACCATATACTCTCTATCCAAAAAACCTGTTGTCAGGAAAAAGCCTAGCAAAGTTTCTTCCTCTCCAT<br>CTTACATTGCGCTAAACATTAATTTAGACACTGACCACTTCAAGATTCA |
| ***PtaHRD* (**populus_717-1B4_scaffolds\|scaffold_22462)<br>CAAAGATTTCCTGTACAGCTAGGTTACGCGGCTAGATCAATTCGTGGTACCTGGGTTATGCTCAGTTGATACGTT<br>AGAAAGATTTTTTAAAAAGAAAAATTAGCAGCAAAATAATACAGAAATGACGAAATAATACAGTTCAGACATGTC<br>ACGATGAATAGATAAAATGAGGTGTTTAAGCAAAAATACCCCATTAGAAACCCTACATGGTAAAAAGAGATGTCT<br>TGCTCAACACAAAAGGCAAGAGCATTAAGAGCACTGTTTAAGATATCATAAGTAATCCTCAGACCATAAAGCTGT<br>CATCTACTGATGATTTATTTTTCTGCCATAAAATATTAATCAAAAGGCAGCAGTAGTACATGAGTAGTCCCACAA<br>ATTTACTGCTGCTAAATTGTCTGCACAAACGTTTGTCAAGTACTTTTTAATGAAACAGGCAGCACTACAACTCAC<br>CCACTGTTTCCCACCATATATATTACAGATAAGGAAATCAAACCCTAAGAAGTGATGTAAGATGTCCAAAAACAT<br>CACTATTCTTGACGTTAAACAAGCTAGTCATGCAGTTTTGCTGCGGAAACCGCAATCCAACCAAGATATCCAACA<br>AAAACACTGGCATACACCCCAACCCTTCACAGACAGCAGTTCATCAACAACTGGTTTGCACTTAAAAATCCAAAC<br>TCAATTAAGATTCTATCTGGAATCCGATCCTCCATAAATGGCTTCTCTCATGGAACACTTGAAGCCATCTTTCCG<br>GCGTCATTCCTTACCTATTATCTTATTGAAAAACCATGCAATATTATCAACAAAGCAACACCTCAAGTGCTGACA<br>GTAGCAGTAGCAGTGGAAGTAGCAGAACTGCTTCTGCTGCTGCTGCTGCTACTGCTGGGGCTTTTGCACCTAAAG<br>TTTCAGGCCACCATCATGTTTACCGTGGAGTCCGGCGTAGGAGTAGTGGAAAATGGGTGTCTGAGATTAGAGA |

***PtaUBQ*** (Populus_717-1B4_scaffolds|scaffold_72584)

AAATATTTTTTTTGAAAAAGCAATACCGCCTCTCACCACCATCCCATTTACAAAATGGAAAGAAACAGAACCATC
CAATCACAAATGTCGACACCTTATTGTTAATTAATTTAATAAAAAAGACGGTCCACCAATTTATAATAATATAAA
GTCCTCACCTTACAGATTCCCTAATTTCACTCTTTCTTTTCTCTTTTTTCTTCTCCCCACCCTCCATTCCTCTCC
CTCTATCTCATCAAACAAATCTCGTAAAAAATTGAAGCCGTTAATTTCCAGAAGGAGAGGAGAGATCCGGTACGT
CTCTCTCGAATTGTGTTCTGGTTTGAGACGATTGATCTGGAGAGAGAGAGAGTTGTCGATGTCGATGTTGATGTT
GATGTTCTTTTTTTTTTTTTTTTGCAGGTTAAGTAGAAGATAAGATGGCATCGAAACGGATCTTGAAGGAATTGAA
AGATTTACAGAAGGATCCTCCTACTTCATGCAGTGCCGGTATTATGACTTTTTTTTTGTTTTTAATTTTTTTCCGA
TTTGTTAAGACTTCCTTTTTAATAGTGTTTTTTTAGTGTTTCGGTTGATCTGTGCCGGTATTATCTTTTTCGTTT
TTTTGGTTGGAATTGATAATGATTTCGTTGATAATATAAGGATGCATGCAGTTGAATAATACTGGCGATGATTAA
CTCTGATCGATAATTTTCTTTGTTAAAATTAAAATTCGATATTTTTAAGGGAAAGCGTAAAACAATATTAATTTT
TTTTTGTTAATTTTTAAAAATTATAGTTGAAAAACTTGTATATTCTGATCACAGGGTGTATTATTTTAGGGTATT
TTTTTTTAAGAAATTATTGTTATCCTGGCTATTCTTAATAATGGTAGGGGGTATCTATGCTGAAT
