## Supplemental Figures S1-S4 for "Temporal modification of H3K9/14ac and H3K4me3 histone marks mediates mechano-responsive gene expression during the accommodation process in poplar"

**Supplementary Figures**

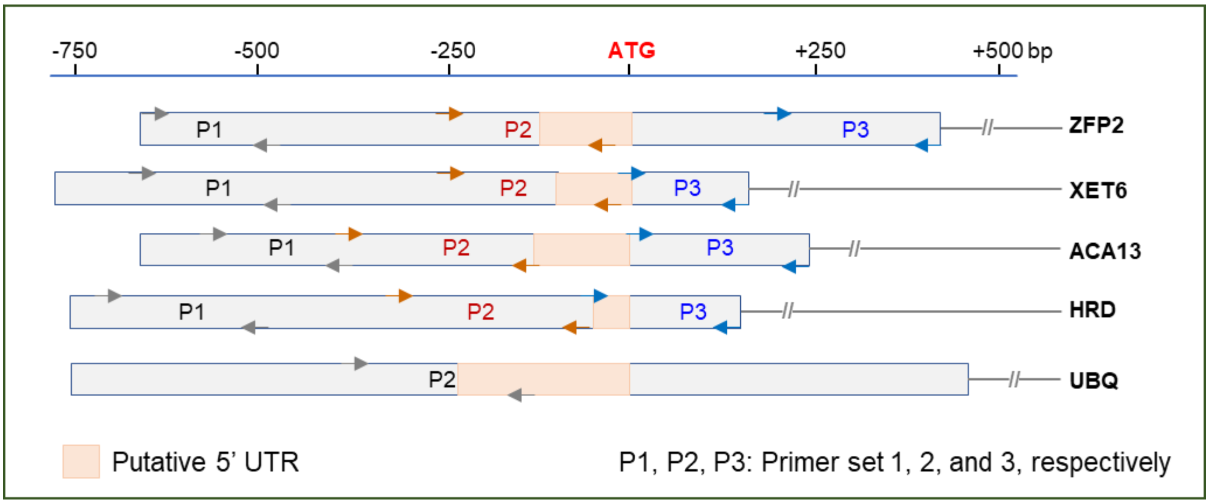


**Figure S1: Schematics show the positions of primers for CHIP-qPCR analysis.** Three sets of primers per gene were designed targeting the following regions: distal (P1) and proximal (P2) regions of the upstream of the start codon, and the first exon (P3). P2 region of *UBQ* was used for normalization. Box in the orange shade is showing the putative 5' UTRs that were identified through pairwise alignment with *P. trichocoharpa* homologs.

| *P. tremula* x *alba* | *P. trichocharpa* homologs | 1B30m vs C30m | 1B2h vs C2h | 1B24h vs C24h | 1B72h vs C72h | 2B30m vs 1B24h | 2B30m vs 1B30m | Description |
|---|---|---|---|---|---|---|---|---|
| ZFP2 | Potri.001G235800 | 55.3 | 3.2 | 1.0 | 1.0 | 10.2 | -5.0 | Zinc finger (C2H2 type) family protein |
| XET6 | Potri.018G094900 | 9.5 | 5.7 | -1.1 | 1.0 | 3.8 | -2.5 | Xyloglucan endotransglucosylase/hydrolase |
| ACA13 | Potri.008G159100 | 3.7 | 2.7 | 1.0 | 1.1 | 1.8 | -2.5 | Potential calcium-transporting ATPase |
| CML11 | Potri.008G159300 | 4.1 | 2.1 | 1.1 | 1.4 | 3.6 | -1.1 | Calcium-binding protein CML |
| RBOHD | Potri.003G159800 | 2.0 | 2.7 | -1.3 | -1.3 | 2.3 | -1.4 | Respiratory burst oxidase homolog protein D |
| FER | Potri.006G110000 | 2.3 | 1.2 | 1.1 | -1.1 | 1.7 | -1.1 | Receptor like protein kinase |
| HRD | Potri.006G021000 | 8.9 | 31.4 | 1.0 | 1.1 | 10.6 | 1.4 | Ethylene-responsive transcription factor |
| JMJ | Potri.003G096100 | 1.7 | 1.2 | 1.0 | 1.0 | 1.4 | -1.3 | Jumonji (Jmj) family protein |
| VIL1 | Potri.018G076500 | 2.5 | 1.9 | 1.0 | 1.0 | 1.8 | -1.3 | VIN3-LIKE PROTEIN 1 |
| HD2A | Potri.009G149400 | 1.1 | 1.5 | 1.8 | 1.1 | 1.0 | 1.4 | Similar to HISTONE DEACETYLASE 2A |
| H1.3 | Potri.007G014200 | -1.4 | -1.3 | -1.7 | -1.3 | -1.3 | -1.7 | Similar to HISTONE H1-3 |
| EZH2 | Potri.014G120100 | -1.3 | -1.3 | -1.7 | 1.0 | 1.2 | -1.4 | Histone-lysine N-methyltransferase EZH2 |
| SDG26 | Potri.005G195400 | -1.1 | -1.1 | -1.7 | 1.1 | 1.4 | -1.3 | SET domain group 26 |
| LHP1 | Potri.019G044400 | n.d. | -1.5 | n.d. | n.d. | n.d. | n.d. | Chromo domain-containing protein |

**Figure S2:** A heat map shows the expression kinetics of mechano-responsive genes determined by microarray analysis (Pomiès *et al.* BMC genomics, 2017) after once or twice mechanostimulation. Poplar plants were subjected to one or two transitory bending treatments with a 24 h interval. The time (0.5, 2, 24, and 72 h) indicates h after 1 bending (1B) or corresponding control (C). Twice mechanostimulated samples (i.e., 2B30m) were harvested 30 min after the second bending. Fold change values were used to draw the heatmap. n.d., not detected.

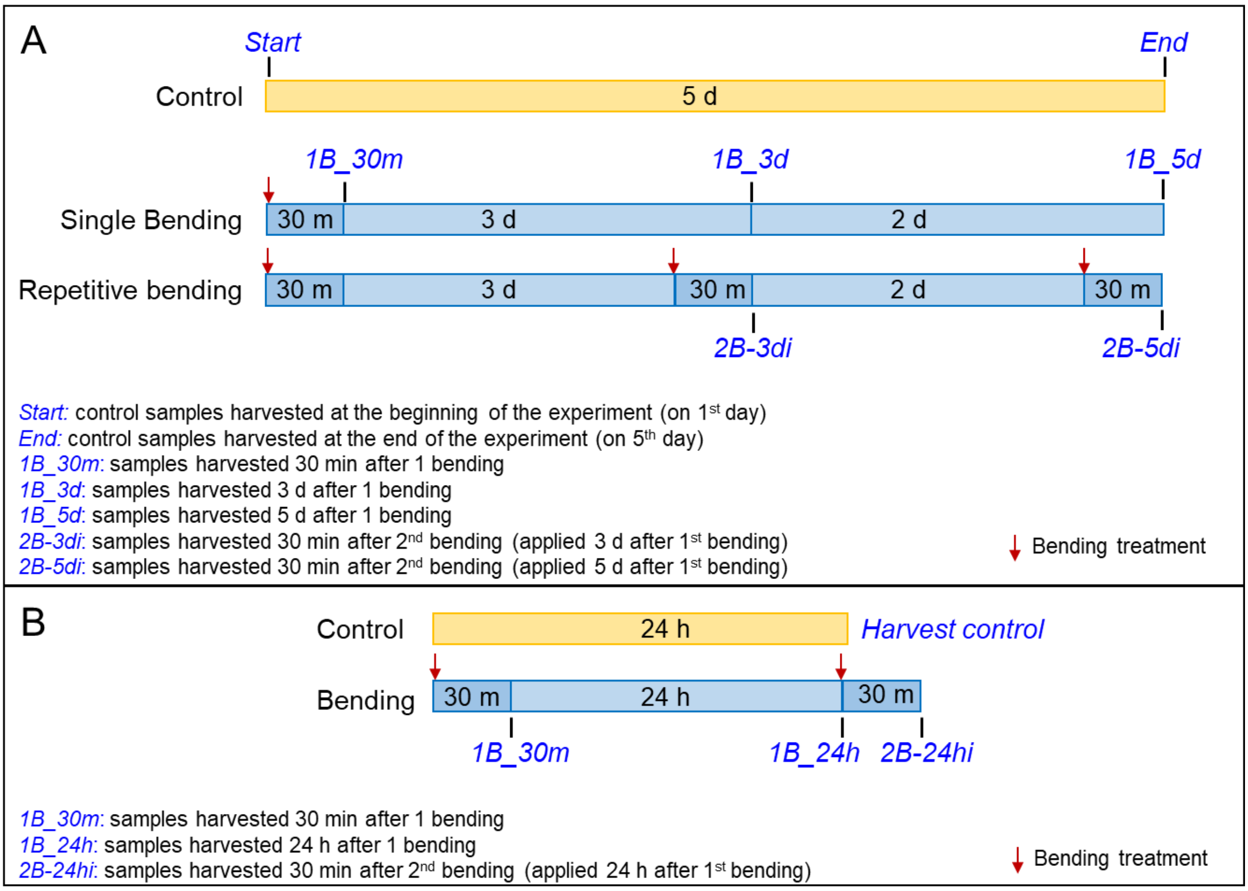


**Figure S3: Schematic representation of experimental strategy.** 1.5 % uniform deformation was applied to 3-month-old poplars by transiently (6 s) bending the basal stem part against a plastic tube as described by Martin *et al* (*J Ex Bot*, 2010). **A)** Poplar plants were subjected to one or two transitory bending treatments at 3- or 5-days intervals. **B)** Poplar plants were subjected to one or two transitory bending treatments with a 24 h interval. Time of bending treatment is indicated with a red arrow.

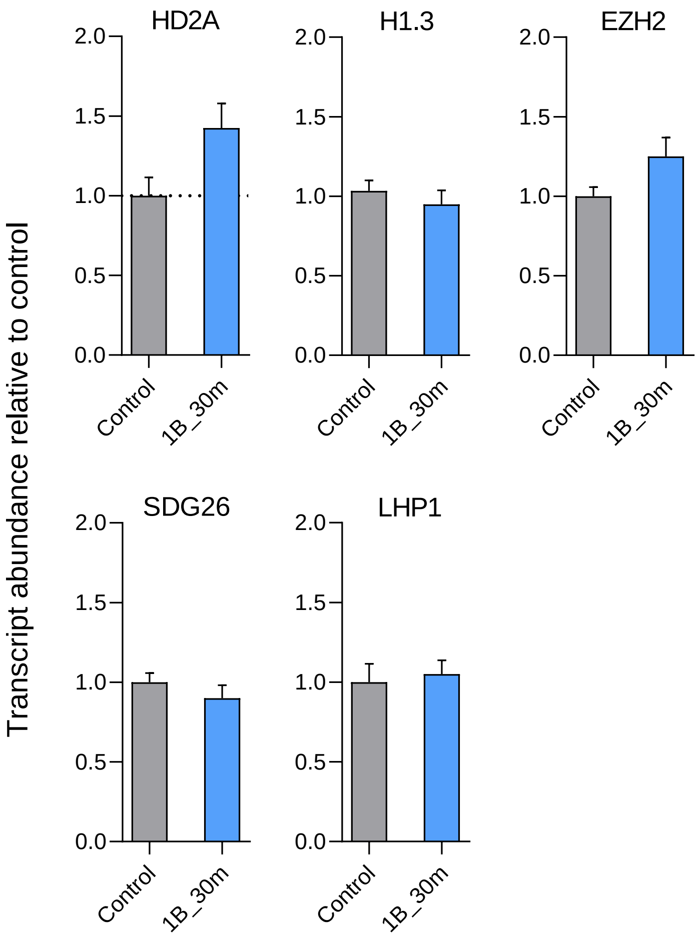


**Figure S4: Relative transcript abundance of *PtaHD2A*, *PtaH1.3*, *PtaEZH2*, *PtaSDG26*, and *PtaLHP1* genes after single transitory stem bending.** Total RNAs were extracted from stems of control (no bent) and 1B_30m (harvested 30 min after 1 bending) plants. The *y* axis denotes transcripts abundance relative to control (set to 1, dotted line) and it was calculated after normalization using 2 reference genes (*UBQ* and *TIP41*) by qPCR. Error bar indicates the standard error of means (*n*=3).
